# MicroRNA signatures of VO_2_peak in older adult participants of the Study of Muscle, Mobility and Aging

**DOI:** 10.1101/2025.01.08.631999

**Authors:** Genesio M. Karere, Fang-Chi Hsu, Russell T. Hepple, Paul M. Coen, Steve Cummings, Anne Newman, Nancy W. Glynn, Lauren Sparks, Nancy E. Lane, Jianzhao Xu, Nathan Wagner, Ge Li, Jeanne Chan, Laura A. Cox, Stephen Kritchevsky

## Abstract

**Background:** Peak oxygen consumption during exercise (VO_2_peak), is a direct measure of cardiorespiratory fitness (CF), a key indicator of physical function and overall health. However, the molecular changes that underpin VO_2_peak variation are not clear. Our objective is to understand the miRNA signatures that relate to VO_2_peak variation, which could provide insights to novel mechanisms that contribute to low VO_2_peak.

**Methods:** We used small RNA sequencing to analyze serum samples from 72 participants (70-79 yrs old, 53% female) of the Study of Muscle, Mobility and Aging (SOMMA). We analyzed samples from individuals with low or high VO_2_peak (N=18/group) as well as samples from 36 randomly selected participants spanning the entire spectrum of VO_2_peak. We used LIMMA analysis package for regression analysis and to identify differentially expressed miRNAs. We used receiver operating characteristic curve analysis to evaluate the Area Under the Curve (AUC) and sensitivity and specificity rates.

**Results:** We identified 1,055 miRNAs expressed in all serum samples. Expression of 65 miRNAs differed between participants with low and high VO_2_peak (p < 0.05). After p-value adjustment, expression of 5 miRNAs (miR-1301-3p, -431-5p, -501-5p, -519a-3p, and -18a-3p) remained significantly different (FDR = 0.05). The five miRNAs had AUC ranging from 0.77 to 0.84. The optimal sensitivity and specificity ranged from 70 to 80% and 80 to 90%, respectively. After adjustment for age and sex covariates, 46 miRNAs significantly correlated with VO_2_peak (p < 0.05) and miR-519a-3p remained significant based on adjusted of p-values.

**Conclusions:** We identified a miRNA signature of VO_2_peak in older individuals that might provide insights to novel mechanisms that drive low VO_2_peak. Future studies will validate the findings in a larger, longitudinal study cohort.

## Introduction

Aging is associated with decreased physical function including impaired mobility (1, 2). The decreased physical function is partially explained by an age-related decline in skeletal muscle mass and strength, or sarcopenia (3, 4). Decline in cardiorespiratory fitness (CRF) is associated with development of sarcopenia, and is highly sensitive to physical fitness differences among older adults (5) and predicts mortality and cardiovascular disease (CVD) (5, 6). Peak oxygen consumption (VO_2_peak), measured during cardiopulmonary exercise testing (CPET), is a direct measure of CF and is positively correlated with lower 400-m walking speed (7), gait strength (8), mitochondrial energetics (9), life-space (5, 10), perceived and performance fatigability (11, 12), and leg power (7, 8). VO2peak also correlates negatively with disease risk, including CVD. Thus, VO_2_peak is a useful indicator of physical function and health status.

Although VO_2_peak is a measure of cardiovascular fitness, indicative of physical function for intervention prescription, the molecular changes that contribute to VO_2_peak variations remain unclear. We posit that microRNAs (miRNAs) are a potential signature of VO_2_peak variations and could provide insights into molecular mechanism driving low VO_2_peak in some older adults. miRNAs are small non-protein coding RNAs that regulate gene expression post-transcriptionally, altering cellular protein abundance. Moreover, the clinical relevance of miRNAs as biomarkers is growing due to their stability and detection in biofluids for diagnosis, prognosis, and responses to treatment, especially exercise interventions (13–16). Despite abundant studies showing that miRNAs are responsive to exercise interventions (13, 17–20) and regulate skeletal muscle development (21–23), there exist little direct proof that blood based miRNAs can discriminate older individuals with low and high VO_2_peak for therapeutic intervention and for screening subjects for clinical trials. A study by Bye et al (24) demonstrated that miR-210, miR-222, and miR-21 are differentially expressed between young adults with low and high VO_2_peak (N=7/group). Other studies showed that plasma miR-23a is associated with percent change in VO_2_peak after exercise in male heart failure patients (25); and that three plasma exosome miRNAs (miR-486-5p, miR-215-5p, miR-941) were upregulated and miR-151b downregulated in response to acute exercise in older men (N=5/group; >65 years old) (26). Importantly, these studies used targeted approaches to analyze expressed serum/plasma miRNAs in small sample sizes. In contrast, we used unbiased small RNA sequencing (small RNA seq) approach to identify all expressed serum miRNAs in older adults and in a larger sample size.

The goal of our baseline study was to identify serum miRNA signatures that accurately discriminate older individuals with low or high VO_2_peak. To identify expressed miRNAs, we leveraged small RNA seq to analyze baseline serum samples from a cohort of participants of the Study of Muscle, Mobility and Aging (SOMMA) (27). We reveal a novel signature of circulating miRNAs for VO_2_peak that correlate with VO_2_peak variation. Our results indicate that blood-based miRNA biomarkers have the potential to identify individuals with low VO_2_peak and provide critical insights into the molecular changes that drive low VO_2_peak.

## Methods

### Study design

We analyzed baseline serum samples from SOMMA participants, including samples from individuals with low or high VO_2_peak (N=18/group), as well as samples from 36 randomly selected participants spanning the entire spectrum of VO_2_peak. The SOMMA was conducted at the University of Pittsburgh (Pittsburgh, PA) and Wake Forest University School of Medicine (Winston-Salem, NC). Inclusion criteria included body mass index (BMI) of 18–40 kg/m^2^; ≥70 years old at enrollment; and eligible for magnetic resonance (MR) imaging and a muscle tissue biopsy. Participants were excluded if unable to walk one quarter of a mile or climb a flight of stairs and having an active malignancy or dementia (27).

### VO_2_peak and sample collections

VO_2_peak (mL/kg/min) was assessed during cardiopulmonary exercise testing (CPET) as previously described (28). Briefly, VO_2_peak was quantified as the highest volume of oxygen consumption averaged over 30-second intervals with progression of exercise intensity in a treadmill, calculated using BreezeSuite metabolic cart software (MGC Diagnostics, St. Paul, MN) and adjusted for participant weight.

### Total RNA isolation

Blood samples were collected at baseline and processed to isolate serum samples following standard protocol. We used 400ul of serum to isolate total RNA using Advanced Serum/Plasma Kit (Qiagen) following the manufacture’s protocol. RNA was quantified using Qubit RNA BR assay (Invitrogen) and DeNovix Fluorimeter (DeNovix).

### Small RNA sequencing

Small RNA seq and sequence analyses were performed as described previously (14, 29). Briefly, we used 20ng of total RNA to generate cDNA libraries using NextFlex Small RNA-Seq Kit v4 (Revvity) and SciClon Workstation Robotics (Revvity). cDNA quality was assessed using TapeStation (Agilent). We performed small RNA seq using Illumina NovaSeq 6000 instrument and reagents. We used mirDeep2 pipeline (30) to analyze fastq-formatted sequence reads with a Phred quality score of 30 or greater, an inferred base call accuracy of 99.9%, to identify miRNA sequences and assess expression levels (read counts). miRNA read counts were normalized holistically using reads per million mapped to a miRNA.

### Data analysis

We used LIMMA (31), a linear model-based R/Bioconductor analysis software with empirical Bayes variance shrinkage to perform differential analysis. Normalized miRNA sequence read counts were log_2_ transformed. Differentially expressed miRNAs between participants with lower (11 to 14.7 mL/kg/min) and higher (28.3-39.4 mL/kg/min) VO_2_peak were identified --- because this was a pilot study, we selected samples from participants with extreme VO_2_peak values to maximize the genetic variability. We also identified expressed miRNAs that correlate with VO_2_peak by performing correlation analysis using 72 samples including the 38 random samples spanning the entire spectrum of VO_2_peak (see Table 1), adjusting for age and sex, where applicable. Further, we performed receiver operating characteristic (ROC) analysis to calculate the Area Under the Curve (AUC), along with sensitivity and specificity, to quantify the predictive ability of each miRNA in classifying the VO_2_peak groups. Additionally, the penalized regression analysis using least absolute shrinkage and selection operator (LASSO) (32), a machine learning approach implemented in the R *glmnet* package, with cross-validation was used to identify miRNAs that predict VO_2_peak. In all analyses, the results were considered significant if the False Discovery Rate (FDR) or adjusted p-value < 0.05.

**Table 1:** Participant characteristics of the SOMMA cohort analyzed in our study.

|  |  |
| --- | --- |
| **Participant characteristics** | **N=72** |
| Age | 76±5.2 |
| Sex: Women | 53% |
| Racial: ethnic minority | 89% White, 8% Black, 3% Asian |
| VO<sub>2</sub>peak (ml/kg/min) | 21.8±8.3 |

## Results

### Study participants

Table 1 shows the characteristics of the study participants. We analyzed samples from 72 participants of SOMMA. Mean age is 76±5.2, 53% females, 89% White, and the average VO_2_peak is 21.8±8.3.

### Sequence reads characteristics

For the sequences that passed the quality filter, the average number of reads per sample was 4,167,566 ranging from 1,336,584 to 8,195,149. The average Phred value for the sequence reads was 35.4 ranging from 32.0 to 36, ensuring that only high-quality sequences were used for statistical analysis.

### Number of expressed miRNAs

We used miRDeep2 tool to analyze miRNA sequences and identified a total of 1,055 miRNAs expressed in serum samples. Table 2 depicts the frequency distribution of expressed miRNAs.

**Table 2:** Distribution of expressed serum miRNAs.

| Read counts | No. of miRNAs |
| --- | --- |
| ≥1,000,000 | 18 |
| ≥100,000 | 55 |
| ≥10,000 | 127 |
| ≥1,000 | 264 |
| ≥500 | 319 |
| ≥100 | 514 |
| ≥10 | 843 |
| ≥5 | 942 |
| ≥1 | 1,055 |

### miRNAs differentially expressed in low and high VO_2_peak

We used LIMMA analysis package and identified 65 miRNAs that differed between individuals with low and high VO_2_peak (Supplemental Table 1) based on nominal p-values. After multiple comparison adjustment using FDR, five miRNAs (miR-1301-3p, miR-431-5p, miR-501-5p, miR-519a-3p, and miR-18a-3p) remained significant (Table 3).

**Table 3:** Topmost significant miRNAs.

| **miRNA ID** | **Log Fold-change** | **p-value** | **FDR<0.05** |
| --- | --- | --- | --- |
| hsa-miR-1301-3p | 2.8 | 0.00008 | 0.02735 |
| hsa-miR-431-5p | 3 | 0.00012 | 0.02735 |
| hsa-miR-501-5p | 2.8 | 0.00018 | 0.02735 |
| hsa-miR-519a-3p | 1.7 | 0.00021 | 0.02735 |
| hsa-miR-18a-3p | 2 | 0.00048 | 0.04936 |

### Predictive values of miRNAs

To identify circulating miRNAs predictive of the VO_2_peak groups, we performed ROC analysis. Interestingly the five miRNAs that showed significant differential expression between low and high VO_2_peak individuals also depicted excellent AUC values ranging from 0.77 to 0.84. The ROC curve analysis revealed high sensitivity and specificity at the optimal threshold point, ranging from 70% to 80% and 80% to 90%, respectively, indicating their potential to discriminate low and high VO_2_peak individuals (Supplemental Figure 1; Table 4)

**Table 4:** Sensitivity and specificity of five serum miRNA biomarkers of VO_2_peak.

| **miRNA ID** | **Sensitivity (%)** | **Specificity (%)** |
| --- | --- | --- |
| hsa-miR-1301-3p | 80 | 90 |
| hsa-miR-431-5p | 70 | 80 |
| hsa-miR-501-5p | 80 | 80 |
| hsa-miR-519a-3p | 70 | 80 |
| hsa-miR-18a-3p | 70 | 80 |

### miRNA correlates of VO_2_peak

Forty-six miRNAs showed significant correlation with VO_2_peak, including miR-1301-3p, miR-431-5p, miR-501-5p, miR-519a-3p, and miR-18a-3p (Supplemental Table 2). After multiple comparison adjustment, miR-519a-3p remained significantly correlated with VO_2_peak (p=0.0001, FDR=0.046 and r=0.5) and miR-18a-3p showed marginal significance and correlation (p=0.0004, FDR=0.092 and r=0.4). Importantly, these two miRNAs are also differentially expressed and show high predictive value to discriminate between individuals with low and high VO_2_peak. We also used machine learning approach and identified miRNAs that predict VO_2_peak, including miR-501-5p, miR-519a-3p, and miR-18a-3p highlighted in Table 5. Together, the findings derived using different approaches confirm the potentiality of these miRNAs as molecular biomarkers of VO_2_peak.

**Table 5:** Serum miRNAs associated with VO_2_peak.

| miRNA ID | Estimate $\beta$ |
| --- | --- |
| hsa-miR-1180-3p | 0.020143 |
| hsa-miR-150-3p | 0.005631 |
| <u>hsa-miR-501-5p</u> | <u>0.27286</u> |
| hsa-miR-1294 | 0.39478 |
| hsa-miR-4665-5p | 0.151921 |
| hsa-miR-517a-3p | 0.793074 |
| hsa-miR-1273h-3p | 0.860643 |
| hsa-miR-323a-5p | 0.251535 |
| <u>hsa-miR-18a-3p</u> | <u>0.880159</u> |
| hsa-miR-106a-5p | 0.002186 |
| hsa-miR-183-3p | 1.249834 |
| hsa-miR-3179 | -0.29502 |
| hsa-miR-1249-3p | 1.020313 |
| <u>hsa-miR-519a-3p</u> | <u>2.26053</u> |
| hsa-miR-1273h-5p | 0.066842 |
| hsa-miR-516b-5p | 1.092905 |
| hsa-miR-6514-5p | 0.3951 |
| hsa-miR-6764-5p | 0.802471 |
| hsa-miR-5010-3p | 0.059 |
| hsa-miR-92a-1-5p | -0.4195 |
| hsa-miR-324-5p | 0.048871 |
| hsa-miR-194-5p | -0.02841 |

## Discussion

The goal of our study was to identify circulating miRNAs as molecular signatures that reliably discriminate older adult participants of SOMMA, with low and high VO_2_peak. We identified a novel panel of five serum circulating miRNAs including miR-1301-3p, miR-431-5p, miR-501-5p, miR-519a-3p, and miR-18a-3p, which were upregulated in individuals with high VO_2_peak compared to those with low VO_2_peak. The expression of the five miRNAs differed significantly between low and high VO_2_peak based on adjusted p-values and showed excellent predictive AUC values with high sensitivity and specificity. Further, the expression of the five miRNAs significantly correlated with VO_2_peak after adjusting for age and sex. miR-519a-3p remained significantly correlated while miR-18a-3p showed marginal correlation after adjusting for age and sex. Together, the signature of the five miRNAs represents a potential miRNA signature that could discriminate individuals with low and high VO_2_peak for clinical interventions and provides insights to molecular changes underpinning low VO_2_peak.

Previous studies have indicated that VO_2_peak is associated with fragility risk factors including mitochondria dysfunction, gait speed, performance fatigability and physical function (7–9, 11, 33, 34). With aging, there is a decline in muscle mass, strength, and aerobic capacity, consistent with lower VO_2_peak. Exercise is an effective intervention strategy to increase CF, skeletal muscle strength and overall physical fitness (35). Importantly, identifying miRNA signatures associated with VO_2_peak in aging populations could also have predictive value in assessing functional capacity, frailty risk, and overall health outcomes. These biomarkers could aid in personalized exercise prescriptions and interventions aimed at improving cardiovascular fitness and maintaining independence in older adults. Moreover, even for older adults tolerant to exercise, monitoring changes in miRNA profiles post-exercise could provide insights into individualized responses to training and potential improvements in VO_2_peak.

Despite the importance of blood-based biomarkers of VO_2_peak, we know little about the potential miRNAs as molecular biomarkers of VO_2_peak. A recent study reported miRNAs associated with VO_2_peak in young (40 to 45 yrs old) individuals (24). The study used a qPCR, a targeted approach, for sample analysis and identified three circulating miRNAs (miR-210, miR-222, and miR-21) that were differentially expressed between individuals with low and high VO_2_peak. A study by Nair *et al.* 2020, identified three plasma exosome miRNAs (miR-486-5p, miR-215-5p, miR-941) were upregulated and miR-151b downregulated in response to acute exercise in older men (N=5/group; >65 years old). Another study by Witvrouwen et al. (25) used qPCR to investigate circulating miRNAs associated with change in VO_2_peak in male patients with heart failure with reduced fraction ejection prescribed exercise intervention for 15 weeks and compared responders to non-responders. The expression of both miR-23a and miR-146a decreased post-exercise. Baseline miR-23a was significantly associated with a percent change in VO_2_peak.

The panel of the five miRNAs we identified represent a potential novel signature of VO_2_peak. These miRNAs are known to play roles in pathogenesis of various human diseases, including cancer, CVD, and Alzheimer’s disease (AD). To our knowledge, no prior study has reported the association of these miRNAs with VO_2_peak. For example, in our study we found that miR-519a-3p was upregulated in high VO_2_peak individuals and correlated significantly with VO_2_peak trait. A prior study revealed that miR-519a-3p is downregulated in glioblastoma and that circNUP98 reduced the inhibitory effects of miR-519-3p on cell proliferation (36). In contrast, other studies showed that the miR-519a-3p was up-regulated exclusively in AD samples from stage I to VI, suggesting its potential use as a novel biomarker of preclinical stages of the disease (37). In cancer studies, miR-519a-3p was overexpressed in more aggressive mutant TP53 breast cancer that was associated with poor survival (38). Serum exosome-derived miR-519a-3p (exo-miR-519a-3p) was significantly upregulated in Gastric Cancer-Liver Metastatic (GC-LM) patients than in patients without LM, indicating a worse prognosis. The exo-miR-519a-3p activates the MAPK/ERK pathway by targeting DUSP2, causing M2-like polarization of macrophages and angiogenesis to augment GC-LM (39). In pregnant women with preeclampsia, miR-519a-3p showed a more than two-fold increase in the blood plasma relative to normal individuals (40). Overall, our findings and literature knowledge show that miR-519a-3p may have diverse effects for different pathophysiological conditions, but its role in VO_2_peak variation and physical function is not known.

The other four serum miRNAs (miR-18a-3p, miR-1301-3P, miR-431-5p and miR-501-5p) are also potential novel biomarkers of VO_2_peak. Previous studies reported that miR-18a-3p was upregulated in patients with immune-mediated necrotizing myopathy and regulates HuR gene (41). miR-1301-3P is upregulated and promotes proliferation and migration in cancer, including esophageal (42), lung (43), ovarian, breast, and endometrial (44). Overexpression of miR-431-5p impairs mitochondrial function in gastric cancer cells (45). This miRNA is also a melanoma tumor and thyroid cancer inhibitor (46). miR-501-5p is a biomarker for essential hypertension (47) and is associated with urinary advanced glycation end products (48). The expression level of miR-501-5p in blood is related to coronary artery disease (49). Together, studies by other investigators showed that these circulating miRNAs play a role in human diseases but none of them have been shown to be associated with VO_2_peak.

## Limitations

An inherent challenge in identifying reliable miRNA biomarkers of VO_2_peak is the analysis of small sample sizes, thus generally underpowered; the use of biassed, targeted approaches; and lack of consideration for gender disparity. Although our study analyzed the largest number of samples so far reported in literature, focused on identifying a signature of circulating miRNA as molecular biomarkers of VO_2_peak in both male and female older adults, the cross-sectional nature of the study limits assessment for generalization, stringent reliability and accuracy of the panel. Hence, there is a need to validate the findings in a larger cohort of SOMMA participants and in longitudinal studies of older adults.

## Conclusion

We have identified a miRNA signature that could discriminate older adults with low or high VO_2_peak and may provide critical insights on the molecular changes that underpin low VO_2_peak variation. While the use of miRNAs as biomarkers for VO_2_peak in aging populations holds promise for understanding physiological aging processes, continued research is needed to elucidate their mechanistic impact on VO_2_peak variation in older adults.

## References

1. Gregg EW, Pereira MA, Caspersen CJ. Physical activity, falls, and fractures among older adults: a review of the epidemiologic evidence. J Am Geriatr Soc. 2000;48(8):883–93.

2. Ryerson B, Tierney EF, Thompson TJ, Engelgau MM, Wang J, Gregg EW, et al. Excess physical limitations among adults with diabetes in the U.S. population, 1997-1999. Diabetes Care. 2003;26(1):206–10.

3. Moreland JD, Richardson JA, Goldsmith CH, Clase CM. Muscle weakness and falls in older adults: a systematic review and meta-analysis. J Am Geriatr Soc. 2004;52(7):1121–9.

4. Yeung SSY, Reijnierse EM, Pham VK, Trappenburg MC, Lim WK, Meskers CGM, et al. Sarcopenia and its association with falls and fractures in older adults: A systematic review and meta-analysis. J Cachexia Sarcopenia Muscle. 2019;10(3):485–500.

5. Newman AB, Kupelian V, Visser M, Simonsick EM, Goodpaster BH, Kritchevsky SB, et al. Strength, but not muscle mass, is associated with mortality in the health, aging and body composition study cohort. The journals of gerontology Series A, Biological sciences and medical sciences. 2006;61(1):72–7.

6. Clausen JSR, Marott JL, Holtermann A, Gyntelberg F, Jensen MT. Midlife Cardiorespiratory Fitness and the Long-Term Risk of Mortality: 46 Years of Follow-Up. Journal of the American College of Cardiology. 2018;72(9):987–95.

7. Ramos SV, Distefano G, Lui LY, Cawthon PM, Kramer P, Sipula IJ, et al. Role of Cardiorespiratory Fitness and Mitochondrial Oxidative Capacity in Reduced Walk Speed of Older Adults With Diabetes. Diabetes. 2024;73(7):1048–57.

8. Healy RD, Smith C, Woessner MN, Levinger I. Relationship between VO2peak, VO2 Recovery Kinetics, and Muscle Function in Older Adults. Gerontology. 2023;69(11):1278–83.

9. Mau T, Lui LY, Distefano G, Kramer PA, Ramos SV, Toledo FGS, et al. Mitochondrial Energetics in Skeletal Muscle Are Associated With Leg Power and Cardiorespiratory Fitness in the Study of Muscle, Mobility and Aging. The journals of gerontology Series A, Biological sciences and medical sciences. 2023;78(8):1367–75.

10. Moored KD, Qiao YS, Rosso AL, Toledo FGS, Cawthon PM, Cummings SR, et al. Dual Roles of Cardiorespiratory Fitness and Fatigability in the Life-Space Mobility of Older Adults: The Study of Muscle, Mobility and Aging (SOMMA). The journals of gerontology Series A, Biological sciences and medical sciences. 2023;78(8):1392–401.

11. Richardson CA, Glynn NW, Ferrucci LG, Mackey DC. Walking energetics, fatigability, and fatigue in older adults: the study of energy and aging pilot. The journals of gerontology Series A, Biological sciences and medical sciences. 2015;70(4):487–94.

12. Qiao YS, Blackwell TL, Cawthon PM, Coen PM, Cummings SR, Distefano G, et al. Associations of accelerometry-measured and self-reported physical activity and sedentary behavior with skeletal muscle energetics: The Study of Muscle, Mobility and Aging (SOMMA). J Sport Health Sci. 2024;13(5):621–30.

13. Juarez M, Castillo-Rodriguez C, Soliman D, Del Rio-Pertuz G, Nugent K. Cardiopulmonary Exercise Testing in Heart Failure. J Cardiovasc Dev Dis. 2024;11(3).

14. Karere GM, Glenn JP, Li G, Konar A, VandeBerg JL, Cox LA. Potential miRNA biomarkers and therapeutic targets for early atherosclerotic lesions. Sci Rep. 2023;13(1):3467.

15. Kim S, Lee WJ, Moon J, Jung KH. Utility of the SERPINC1 Gene Test in Ischemic Stroke Patients With Antithrombin Deficiency. Front Neurol. 2022;13:841934.

16. Peng F, Li H, Xiao H, Li L, Li Y, Wu Y. Identification of a three miRNA signature as a novel potential prognostic biomarker in patients with bladder cancer. Oncotarget. 2017;8(62):105553–60.

17. Afzal M, Greco F, Quinzi F, Scionti F, Maurotti S, Montalcini T, et al. The Effect of Physical Activity/Exercise on miRNA Expression and Function in Non-Communicable Diseases-A Systematic Review. Int J Mol Sci. 2024;25(13).

18. Al-Rawaf HA, Gabr SA, Iqbal A, Alghadir AH. Circulating microRNAs as potential biomarkers of physical activity in geriatric patients with HCV. BMC Mol Cell Biol. 2024;25(1):18.

19. Huang LY, Lim AY, Hsu CC, Tsai YF, Fu TC, Shyu YC, et al. Sustainability of exercise-induced benefits on circulating MicroRNAs and physical fitness in community-dwelling older adults: a randomized controlled trial with follow up. BMC Geriatr. 2024;24(1):473.

20. Tee CCL, Chong MC, Cooke MB, Rahmat N, Yeo WK, Camera DM. Effects of exercise modality combined with moderate hypoxia on blood glucose regulation in adults with overweight. Front Physiol. 2024;15:1396108.

21. Artigas-Arias M, Curi R, Marzuca-Nassr GN. Myogenic microRNAs as Therapeutic Targets for Skeletal Muscle Mass Wasting in Breast Cancer Models. Int J Mol Sci. 2024;25(12).

22. Kielbowski K, Bakinowska E, Procyk G, Zietara M, Pawlik A. The Role of MicroRNA in the Pathogenesis of Duchenne Muscular Dystrophy. Int J Mol Sci. 2024;25(11).

23. O’Bryan SM, Lavin KM, Graham ZA, Drummer DJ, Tuggle SC, Van Keuren-Jensen K, et al. Muscle-Derived microRNAs Correlated with Thigh Lean Mass Gains during Progressive Resistance Training in Older Adults. J Appl Physiol (1985). 2024.

24. Bye A, Rosjo H, Aspenes ST, Condorelli G, Omland T, Wisloff U. Circulating microRNAs and aerobic fitness--the HUNT-Study. PloS one. 2013;8(2):e57496.

25. Witvrouwen I, Gevaert AB, Possemiers N, Beckers PJ, Vorlat A, Heidbuchel H, et al. Circulating microRNA as predictors for exercise response in heart failure with reduced ejection fraction. Eur J Prev Cardiol. 2021.

26. Nair VD, Ge Y, Li S, Pincas H, Jain N, Seenarine N, et al. Sedentary and Trained Older Men Have Distinct Circulating Exosomal microRNA Profiles at Baseline and in Response to Acute Exercise. Front Physiol. 2020;11:605.

27. Cummings SR, Newman AB, Coen PM, Hepple RT, Collins R, Kennedy Ms K, et al. The Study of Muscle, Mobility and Aging (SOMMA): A Unique Cohort Study About the Cellular Biology of Aging and Age-related Loss of Mobility. The journals of gerontology Series A, Biological sciences and medical sciences. 2023;78(11):2083–93.

28. Wolf C, Blackwell TL, Johnson E, Glynn NW, Nicklas B, Kritchevsky SB, et al. Cardiopulmonary Exercise Testing in a Prospective Multicenter Cohort of Older Adults: The Study of Muscle, Mobility and Aging (SOMMA). medRxiv. 2023.

29. Karere GM, Cox LA, Bishop AC, South AM, Shaltout HA, Mercado-Deane MG, et al. Sex Differences in MicroRNA Expression and Cardiometabolic Risk Factors in Hispanic Adolescents with Obesity. J Pediatr. 2021;235:138–43 e5.

30. Friedlander MR, Mackowiak SD, Li N, Chen W, Rajewsky N. miRDeep2 accurately identifies known and hundreds of novel microRNA genes in seven animal clades. Nucleic acids research. 2012;40(1):37–52.

31. Ritchie ME, Phipson B, Wu D, Hu Y, Law CW, Shi W, et al. limma powers differential expression analyses for RNA-sequencing and microarray studies. Nucleic acids research. 2015;43(7):e47.

32. Tibshirani R. The lasso method for variable selection in the Cox model. Stat Med. 1997;16(4):385–95.

33. 33. Qiao YS, Blackwell TL, Cawthon PM, Coen PM, Cummings SR, Distefano G, et al. Associations of Objective and Self-Reported Physical Activity and Sedentary Behavior with Skeletal Muscle Energetics: The Study of Muscle, Mobility and Aging (SOMMA). medRxiv. 2023.

34. Vasquez-Gomez J, Cifuentes-Amigo A, Castillo-Retamal M, Zamuner AR. A VO(2peak) prediction model in older adults’ patients with Parkinson’s disease. Exp Gerontol. 2023;181:112285.

35. Pahor M. Randomized controlled trials involving multidisciplinary interventions in the community. The journals of gerontology Series A, Biological sciences and medical sciences. 2006;61(5):472–3.

36. Lu J, Lou G, Jiang L, Liu X, Jiang J, Wang X. CircNUP98 Suppresses the Maturation of miR-519a-3p in Glioblastoma. Front Neurol. 2021;12:679745.

37. Jacome D, Cotrufo T, Andres-Benito P, Lidon L, Marti E, Ferrer I, et al. miR-519a-3p, found to regulate cellular prion protein during Alzheimer’s disease pathogenesis, as a biomarker of asymptomatic stages. Biochim Biophys Acta Mol Basis Dis. 2024;1870(5):167187.

38. Breunig C, Pahl J, Kublbeck M, Miller M, Antonelli D, Erdem N, et al. MicroRNA-519a-3p mediates apoptosis resistance in breast cancer cells and their escape from recognition by natural killer cells. Cell Death Dis. 2017;8(8):e2973.

39. Qiu S, Xie L, Lu C, Gu C, Xia Y, Lv J, et al. Gastric cancer-derived exosomal miR-519a-3p promotes liver metastasis by inducing intrahepatic M2-like macrophage-mediated angiogenesis. J Exp Clin Cancer Res. 2022;41(1):296.

40. Timofeeva AV, Gusar VA, Kan NE, Prozorovskaya KN, Karapetyan AO, Bayev OR, et al. Identification of potential early biomarkers of preeclampsia. Placenta. 2018;61:61–71.

41. Ye L, Zuo Y, Chen F, Peng Q, Lu X, Wang G, et al. miR-18a-3p and Its Target Protein HuR May Regulate Myogenic Differentiation in Immune-Mediated Necrotizing Myopathy. Front Immunol. 2021;12:780237.

42. Du J, Zhang S, Zhang X, Yang Z, Xue S, Xu G, et al. miR-1301-3p promotes invasion and migration and EMT progression in esophageal cancer by downregulating NBL1 expression. Thorac Cancer. 2023;14(30):3032–41.

43. Qu M, Jin Z, Xu Y, Sun W, Luo Y, Zhang N, et al. hsa-miR-1301-3p Promotes the Proliferation and Migration of Nonsmall Cell Lung Cancer Cells and Reduces Radiosensitivity via Targeting Homeodomain-Only Protein Homeobox. Genet Test Mol Biomarkers. 2023;27(12):393–405.

44. Pane K, Zanfardino M, Grimaldi AM, Baldassarre G, Salvatore M, Incoronato M, et al. Discovering Common miRNA Signatures Underlying Female-Specific Cancers via a Machine Learning Approach Driven by the Cancer Hallmark ERBB. Biomedicines. 2022;10(6).

45. Wu J, Deng Z, Zhu Y, Dou G, Li J, Huang L. [Overexpression of miR-431-5p impairs mitochondrial function and induces apoptosis in gastric cancer cells via the Bax/Bcl-2/caspase3 pathway]. Nan Fang Yi Ke Da Xue Xue Bao. 2023;43(4):537–43.

46. Gao Y, Wang Y, Xu L, Xie X, Zhu L, Wang F. CircRTN1 acts as a miR-431-5p sponge to promote thyroid cancer progression by upregulating TGFA. Hormones (Athens). 2022;21(4):611–23.

47. Jusic A, Junuzovic I, Hujdurovic A, Zhang L, Vausort M, Devaux Y. A Machine Learning Model Based on microRNAs for the Diagnosis of Essential Hypertension. Noncoding RNA. 2023;9(6).

48. Russo P, Lauria F, Sirangelo I, Siani A, Iacomino G. Association between Urinary AGEs and Circulating miRNAs in Children and Adolescents with Overweight and Obesity from the Italian I.Family Cohort: A Pilot Study. J Clin Med. 2023;12(16).

49. Liu L, Zhang J, Wu M, Xu H. Identification of key miRNAs and mRNAs related to coronary artery disease by meta-analysis. BMC Cardiovasc Disord. 2021;21(1):443.

